# PolyRNN: A time-resolved model of polyphonic musical expectations aligned with human brain responses

**DOI:** 10.1101/2024.11.27.625704

**Authors:** Paul Robert, Mathieu Pham Van Cang, Manuel Mercier, Agnès Trébuchon, Fabrice Bartolomei, Luc H. Arnal, Benjamin Morillon, Keith Doelling

## Abstract

Musical expectations shape how we perceive and process music, yet current computational models are limited to monophonic or simplified stimuli. The study of the neural processes underlying musical expectations in real-world music therefore requires significant advances in our statistical modeling of these stimuli. We present PolyRNN, a recurrent neural network designed to model expectations in naturalistic, polyphonic music. We recorded neurophysiological activity non invasively (MEG) and within the human brain (intracranial EEG) while participants listened to naturally expressive piano recordings. The musical expectations estimated by the model are encoded in evoked P2- and P3-like components in auditory regions. Comparing PolyRNN to a state-of-the-art generative music model, we show that piano roll representations are best suited to represent expectations in polyphonic contexts. Overall, our approach provides a new way to capture the musical expectations emerging from natural music listening, and enables the study of predictive processes in more ecologically valid settings.

## Introduction

Music is more than sound. It is composed of organized patterns, shaped intentionally by musicians to convey meaning, evoke emotions, and move people to dance. A large body of research has shown that listeners form expectations about musical patterns such as melodies, rhythms, or harmony. When these expectations are fulfilled or violated, they exhibit measurable behavioral and neurophysiological responses, in line with the predictive coding framework (Koelsch et al., 2019). This phenomenon has been extensively studied in the 1980s and 1990s with reductionist experimental paradigms, which relied on artificial stimuli to manipulate the predictability of individual musical events (Krumhansl & Shepard, 1979; Krumhansl & Kessler, 1982; Bharucha & Stoeckig, 1986; 1987; Janata 1995; Besson & Faïta, 1995; Patel et al., 1998; Koelsch et al., 2000). In the following years, several computational models were proposed to represent and quantify the listener’s expectations (Tillmann et al., 2000; Temperley, 2008; 2009; Pearce, 2005; 2018; Verosky & Morgan, 2021). In parallel, psychology and neuroscience of music have moved towards more naturalistic stimuli, in an effort to target the social and affective mechanisms at play in real-world music listening (Mas-Herrero et al., 2012; 2018; Tervaniemi, 2023; Abrams et al., 2024; Curzel et al., 2024). Beyond their theoretical significance, computational models offer a practical tool to evaluate musical expectations in real music samples, and enable us to study the relationship between perceptual predictions and other cognitive processes (Egermann et al., 2013; Cheung et al., 2019; 2024; Gold et al., 2019; 2023; Zioga et al., 2020; Bonetti et al., 2024).

Models of musical expectations aim at simulating two cognitive mechanisms (Pearce, 2005; 2018). First, *statistical learning* refers to the listener’s ability to acquire knowledge about musical structures through regular exposure – even without explicit training. All current computational models rely on machine learning algorithms that extract regularities from a dataset, and the choice of musical corpus and training method define how the learning process is approximated. For instance, the Information Dynamics of Music (IDyOM; Pearce, 2005; 2018) model provides an explicit implementation of statistical learning through Markov models (*i.e.*, frequency counts). More recent deep learning models like MelodyRNN (Sankaran et al., 2024) or the Music Transformer (Huang et al., 2018; Kern et al., 2022) rely on training algorithms optimized for neural networks (*i.e.*, loss functions, gradient descent and optimizers), and offer a less interpretable learning method. Second, *contextual predictions* represent the events that a listener expects based on the immediate context and prior learning. To simulate this process, computational models are designed to yield quantitative estimates of the probability of each possible event given the context, and measures of surprise (prediction error) and uncertainty (precision of predictions) are typically used to predict behavioral and neurophysiological responses to music (Pearce, 2018; Di Liberto et al., 2020; Kern et al., 2022; Sankaran et al., 2024).

One key limitation of current models of music prediction used to model human neural responses is that they only process single-stream sequences (i.e. sequences of events unfolding one by one), whereas natural music is predominantly polyphonic (i.e. multiple notes can be played at once). With IDyOM for instance, this constraint arises from the use of Markov models at its core, and restricts its learning and predictions to melodies or chord series. In deep learning models, the choice of musical representation dictates what is being predicted, so that models can be tailored for processing melodies (MelodyRNN) or polyphonic music (Music Transformer). Still, previous studies using these neural networks relied on monophonic stimuli only (Kern et al., 2022; Sankaran et al., 2024), and their ability to predict behavior or neural responses in polyphonic context remains to be tested. Alternatively, expectations in polyphony have been handled with Markov models (D-REX; Skerritt-Davis et al., 2019) operating on acoustic descriptors (Abrams et al., 2024). However, this approach does not address the patterns emerging from discrete events like notes and chords, as other models do, nor is it trained to learn the symbolic relationships of specific styles of music. Overall, the lack of computational model of expectations in polyphony prevents the use of more naturalistic music in experiments. Since most popular music is polyphonic, the mismatch between single-stream stimuli and real music substantially reduces ecological validity. Moreover, current models are unable to account for the knowledge of polyphonic music that people may have developed through exposure, such as the interplay between a bass line and a melody for instance.

With the advent of deep learning in recent years, a wide range of models have been developed for music generation (Civit et al., 2022). In particular, recurrent neural networks (RNN) and Transformers for symbolic music follow a synthesis process that includes estimating the probability of upcoming notes in a sequence, and can thus be used as models of musical expectations. Some of these models, like Performance RNN (PerfRNN; Oore et al., 2020), have been specifically developed to generate expressive polyphonic music, making them promising candidates for modeling expectations in naturalistic stimuli. One key consideration, however, is the musical representation on which such models operate. To ensure computational efficiency and scalability, the musical sequences are transformed into midi tokens, coding for instance the onset of a note or the time between two events (Fradet et al., 2023). In polyphonic contexts, this results in a representational format that is unlikely to reflect the expectations of a human listener. For instance, when multiple notes are played simultaneously, each note is encoded as a separate token and predicted sequentially, disrupting the temporal relationship between events. The main alternative to midi tokens is the piano-roll representation, in which events are placed in a 2D matrix with time and pitch as dimensions. This preserves the relationship between events and may better reflect how listeners perceive and form expectations about polyphonic music. In practice, existing piano-roll-based models like MidiNet (Yang & Yang, 2017) or MuseGAN (Dong et al., 2018) are designed to generate entire chunks of music at once. While efficient for music generation, a model of human expectations must be able to estimate the occurrence probability of possible events at each time step of a musical sequence. Overall, deep learning models offer a promising avenue for capturing the complex structure of polyphonic music. Specifically, using sequential networks like RNNs combined with a piano-roll representation could provide a plausible computational model of the listeners’ musical expectations.

In this paper, we introduce and validate a model of musical expectations for naturalistic, polyphonic music. We trained a recurrent neural network, PolyRNN, on a large dataset of real piano performances (classical music) to perform a multi-item prediction task with a piano-roll representation of the recordings (Figure 1A; see Video S1). The model yields a probability of occurrence for each possible note at each timestep of the piano roll. From these predictions, we extract three key metrics: (i) surprise (prediction error), reflecting the difference between predicted and actual note occurrences; (ii) uncertainty (entropy), reflecting the uncertainty among multiple possibilities; and (iii) predicted density, indicating the likelihood of note events occurring (Figure 1C). We first verified the musical capabilities of the model by comparing its surprise values on the musical genre it was trained on (classical) versus novel genres it was not exposed to during training (jazz, pop). Second, taking IDyOM as an established benchmark, we compared the encoding of PolyRNN’s or IDyOM’s features on an openly-available EEG dataset acquired with monophonic pieces (Di Liberto et al., 2020). Then, we acquired intracranial EEG (sEEG) and MEG data while participants listened to natural, polyphonic piano recordings, and we tested whether PolyRNN’s predictive metrics align with neural responses to music, to test whether they capture aspects of human auditory expectations. Finally, to evaluate the impact of the model’s input representation, we compared PolyRNN (piano-roll) and PerfRNN (midi token) on the same sEEG and MEG data.

**Figure 1.**
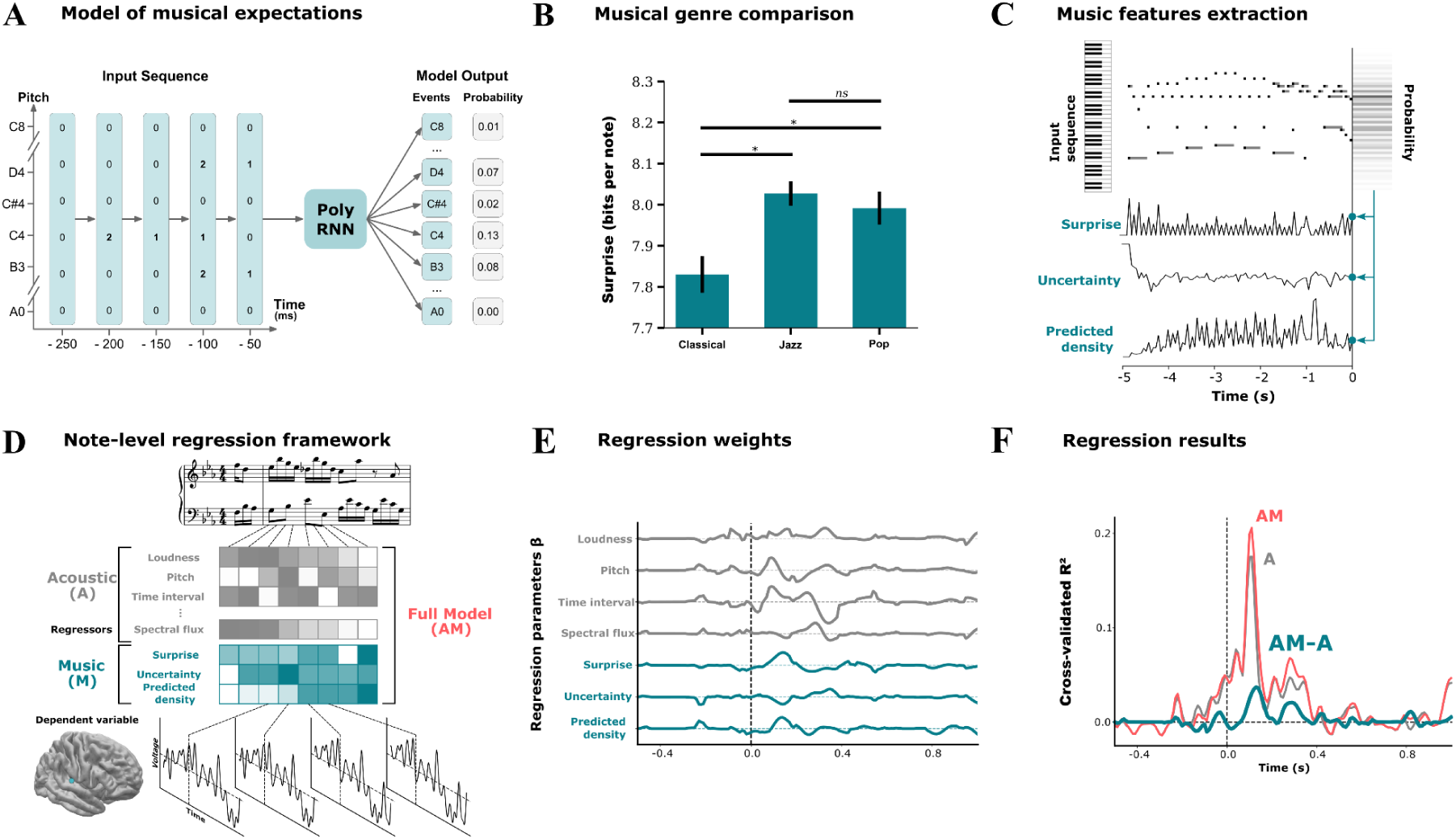
PolyRNN working principles and analysis pipeline. **(A)** Description of PolyRNN working principles. PolyRNN receives sequences of vectors coding for the start (2), the sustain (1) or the absence (0) of each possible note of the 88 notes of the piano keyboard, with one vector per timestep (50 ms). With this representation, multiple note onsets can be represented within a single timestep. The model outputs predictions about the next timestep with one probability of occurrence per note. **(B)** Average surprise (prediction error) per note of PolyRNN on unseen musical corpora of various genres, after the training on a large corpus of classical piano music. *: p < 0.05 for paired t-tests with Bonferroni correction. Error bars indicate SEM across music samples. **(C)** In experimental stimuli, three metrics are extracted from PolyRNN’s predictions at each timestep: Surprise, Uncertainty and Predicted density. **(D) Top:** Each piano note in the experiment is associated with a series of acoustic features (gray; up to k=29) and with the metrics derived from the models’ outputs (cyan). **Bottom:** Neural signals are epoched around note onsets. Epochs of an exemplar sEEG channel (right auditory cortex) are shown. **(E)** Five-fold cross-validated ridge regression to predict the neural response to individual notes. Beta coefficients of a subset of acoustic and music model features are shown, obtained on the exemplar sEEG channel from (C). **(F)** Variance explained by all (acoustic and model) features (AM) in the exemplar sEEG channel, acoustic features only (A), and uniquely explained by model features (ΔR² = AM - A).

## Results

### 1. PolyRNN captures high-order musical features

We first compared the surprise of PolyRNN on unseen musical corpora with musical genres that either matched or differed with the training corpus. The model has a lower average surprise per note in classical music compared to jazz (t(908.7) = 3.7, p < 0.001) or pop music (t(791.1) = 2.7, p = 0.02; Figure 1B). This indicates that during the training, PolyRNN captured genre-specific patterns, possibly corresponding to higher-order musical features. We then compared PolyRNN and the well-established IDyOM model in their ability to predict brain responses to music. As IDyOM can only process single stream sequences, we tested both models using an existing open-source set of electroencephalography (EEG) data acquired during monophonic music listening (Di Liberto et al., 2020). We predicted the amplitude of the neural signal in response to individual notes (Figure 1D-F) using basic acoustic features – temporal envelope and derivative – only (A), or using a combination of acoustic and IDyOM’s music features (AM). The difference between these models (ΔR² = AM - A) shows that the music features extracted from IDyOM predict a P2-like neural response over central electrodes around 200 ms (unilateral t-test vs. zero: t(19) = 4.4, p < 0.001; Figure 2A; see Di Liberto et al., 2020). PolyRNN better predicts this component of neural response to music notes (unilateral t-test vs. zero: t(19) = 6.3, p < 0.001; PolyRNN vs. IDyOM bilateral paired t-test: t(19) = 5.2, p < 0.001; Figure 2B). To investigate whether this effect can be explained away by more complex features, such as distributions of note pitch and timing, we replicated these analyses with additional regressors (see Method). While IDyOM’s explained variance can be accounted for by these low-order contextual features (unilateral t-test vs. zero: t(19) = -4.5, p = 1), PolyRNN captures additional higher-order brain activity (unilateral t-test vs. zero: t(19) = 3.2, p = 0.002; PolyRNN vs. IDyOM bilateral paired t-test: t(19) = 4.52, p < 0.001; see Fig. S1 for additional analyses).

**Figure 2.**
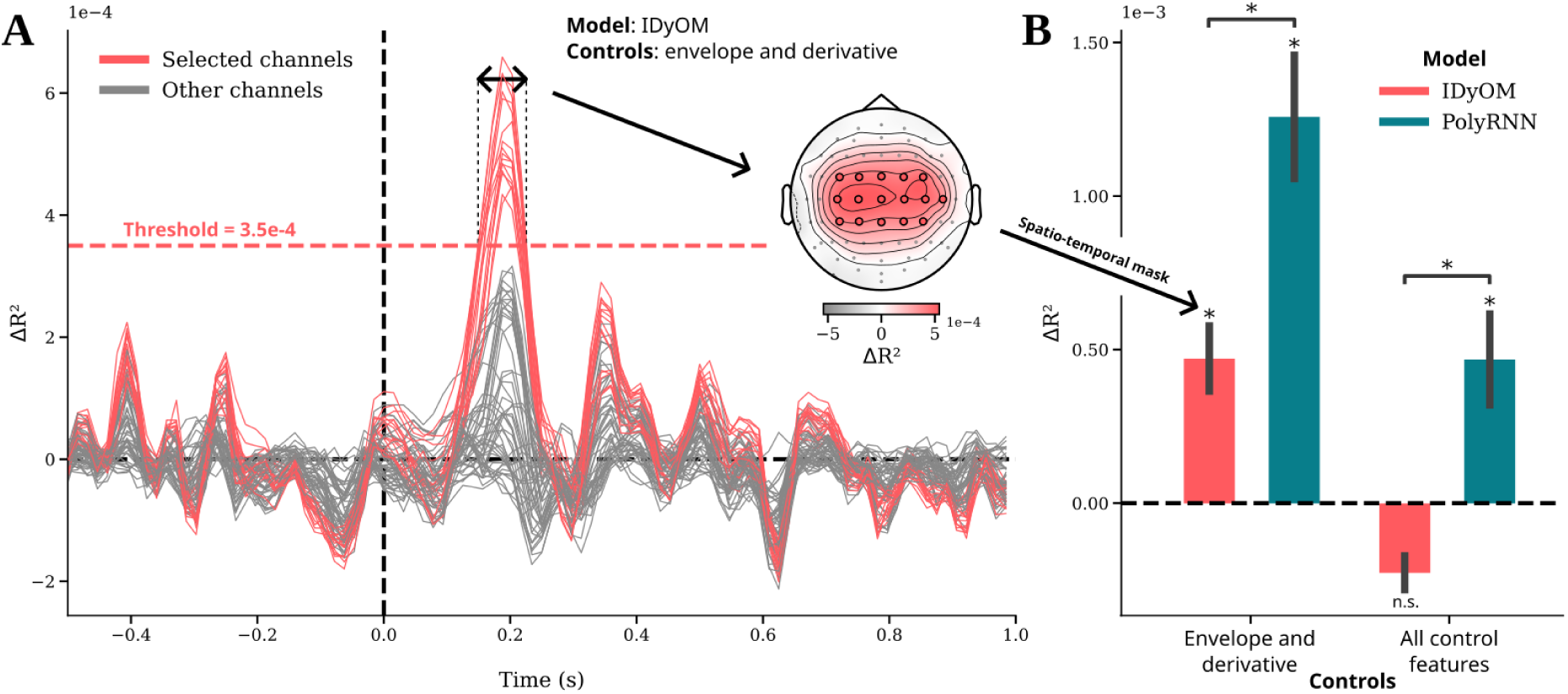
Comparison of PolyRNN and IDyOM models with monophonic music in EEG. **(A)** Replication of the results of Di Liberto and colleagues (2020). Plotted lines represent the variance uniquely explained by IDyOM’s features (ΔR² = AM - A; see Fig. 1F) at the note-level (note onsets locked at t = 0s) for each EEG channel, averaged across participants. The sound envelope and its half-way rectified derivative were used as control acoustic features. To capture the peak effect, all channels and timepoints with ΔR² values exceeding an arbitrary threshold of 3.5*10^-4^ are kept as a cluster of interest for subsequent models comparison. **(B)** Variance uniquely explained by IDyOM and PolyRNN features, averaged within the spatiotemporal mask. The analysis was conducted by controlling for the variance captured either by (left) the acoustic temporal envelope and its derivative alone (as in Di Liberto and colleagues, 2020), or (right) with additional descriptors of note pitch and timing (see Methods). *: p < 0.05 for paired t-tests with Bonferroni correction. Error bars indicate SEM across participants (N = 20).

### 2. Polyphonic expectations are encoded in the cortical response to music

Next, we investigated the encoding of polyphonic expectations estimated by PolyRNN in the human brain. We recorded neural activity from 27 healthy participants in magnetoencephalography (MEG) and 10 patients in stereotactic EEG (sEEG) as they listened to recordings of ecological classical piano pieces. We predicted the amplitude of the neural signal in response to individual or simultaneous notes (Figure 1C) with acoustic only (A; see Figure S2, S3) or acoustic and PolyRNN’s model features (AM; see Method).

In sEEG, PolyRNN predicts (ΔR² = AM - A) a P2-like neural response (61 significant temporal clusters encompassing 8/10 patients; cluster-based permutation test, ΔR²-threshold = 0.005, p < 0.05), mostly in primary and associative auditory regions (Figure 3A, 3C). In MEG, PolyRNN features are encoded in temporal sensors around 200 ms after note onset (12 significant spatiotemporal cluster; cluster-based permutation test, t-threshold = 2.63, p < 0.05; Figure 3B, 3F). No difference was observed between left and right hemisphere clusters (t-threshold = 2.06, p > 0.9, two-tailed; Figure S4). To confirm these effects with an independent channel selection, we measured the improvement in predicted variance (ΔR² = AM - A) on the channels with the highest auditory response per participant (see Method). PolyRNN significantly improves the explained variance in those channels in both sEEG (in 10/10 patients; paired t-test, t(9) = 2.91, p = 0.017, two-tailed; Figure 3D) and MEG (in 22/27 participants; paired t-test, t(26) = 3.44, p < 0.002, two-tailed; Figure 3G).

**Figure 3.**
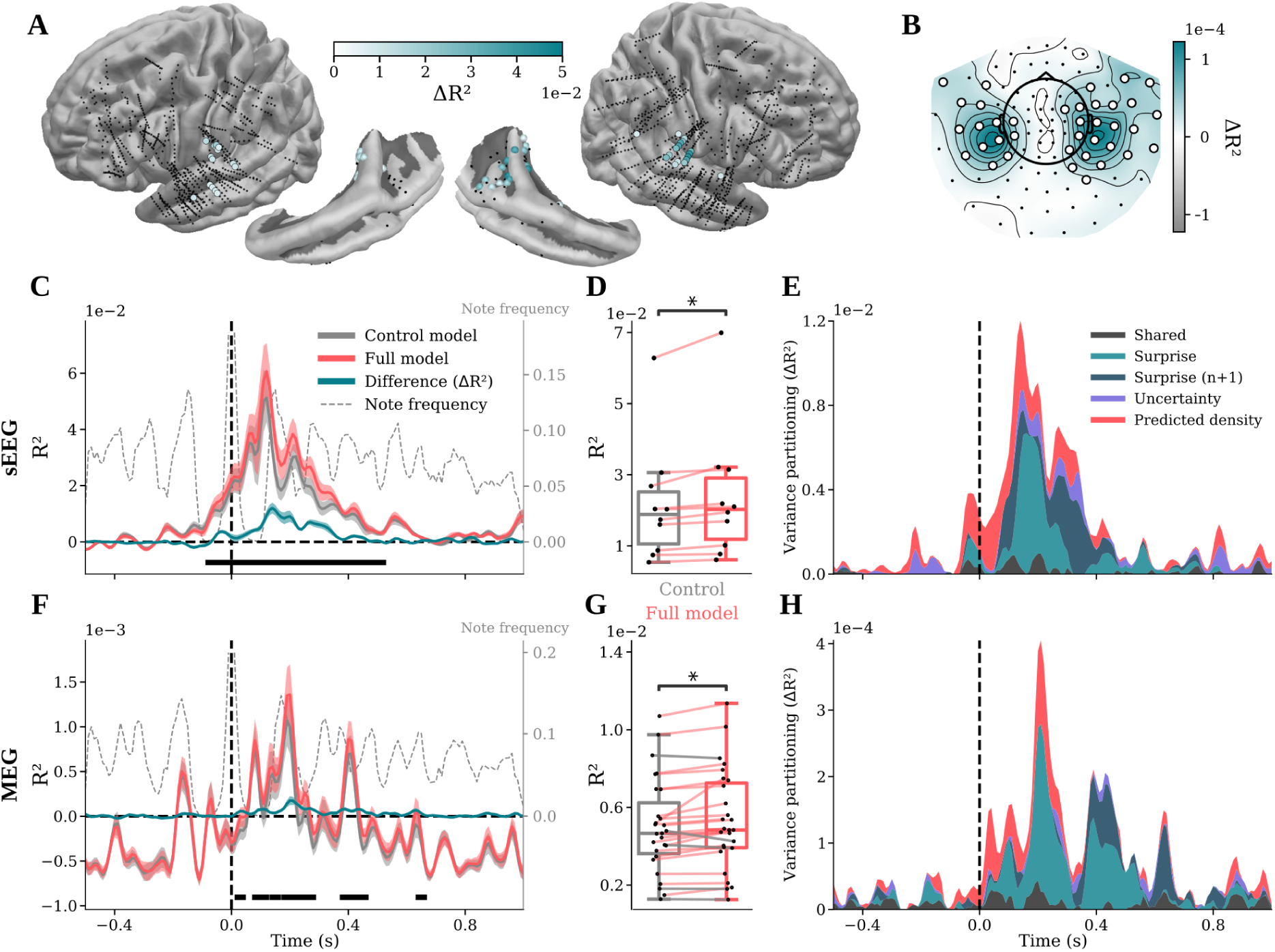
Encoding of polyphonic expectations (PolyRNN) in sEEG and MEG. **(A)** Variance uniquely explained by PolyRNN in sEEG data (N = 10), plotted on whole brains and temporal regions. Peak ΔR² (within [-0.5, 1] s) are color coded in significant sEEG channels. Black dots indicate non-significant channels (N = 10 patients). **(B)** Topography of ΔR² values averaged across time ([-0.5, 1] s) and participants in MEG data (N = 27). Significant channels are indicated as white dots (p < 0.05, cluster corrected). **(C)** Variance explained by PolyRNN and/or control acoustic features at the note-level (note onsets locked at t = 0s) in sEEG, averaged across participants and significant channels. Shaded areas are SEM across channels. Solid black ribbon indicate significant differences between control and full models in at least one spatio-temporal cluster (p < 0.05, cluster corrected). Note frequency represents the frequency of occurrence of a note at each timepoint (p = 1 at t = 0s, right y-axis). **(D)** Individual R² peaks (within [-0.5, 1] s) averaged across auditory channels (see Method) in sEEG. Colored lines indicate individual performance increase uniquely provided by PolyRNN music features. Error bars indicate SEM across participants. **(E)** Cumulative plot of variance explained, partitioned by each PolyRNN music feature, in sEEG. Surprise (n+1) corresponds to the surprise of the next note. The shared variance represents all non-unique explained variances. The Y-axis is depicted from zero as negative cross-validated R2 is meaningless. **(F-G-H)** Same as **(C-D-E)** with MEG data. **(F)** Shaded areas are SEM across participants.

Finally, we used variance partitioning (Figure 3E, 3H) to disentangle the contribution of the different PolyRNN music features, namely surprise, uncertainty and predicted density (see Method). In both MEG and sEEG, most of the effect is driven by the surprise of the current note. The encoding of predicted density arises earlier than surprise, while the encoding of uncertainty remains low overall. Lastly, in MEG, the surprise of the current note is associated with a P3-like neural response, which extends into the processing of the next note (n+1; >350 ms; Figure 3H).

### 3. Musical expectations are best represented with piano-roll based modeling

To evaluate the impact of the model’s input representation, we compared the ability of PolyRNN and PerfRNN to predict the brain’s response to polyphonic music. These two neural networks have similar architecture (recurrent units) and have been trained on the same musical corpus, but PolyRNN is tasked to predict the content of the next timestep in a piano roll while PerfRNN predicts the next midi token in a sequence. In sEEG, PolyRNN outperforms PerfRNN in predicting the P2-like neural response, principally along the temporal lobe (Figure 4A-B). On the channels with the highest auditory response (see Method), PolyRNN better predicts the brain’s response to music than PerfRNN (t(199) = 7.89, p < 0.001; Figure 4C). In MEG, we obtained similar results, with the highest difference between models around 200ms after note onset in temporal and frontal sensors (4 significant spatiotemporal clusters; cluster-based permutation test, t-threshold = 2.53, p < 0.05, two-tailed; Figure 4D-E).

**Figure 4.**
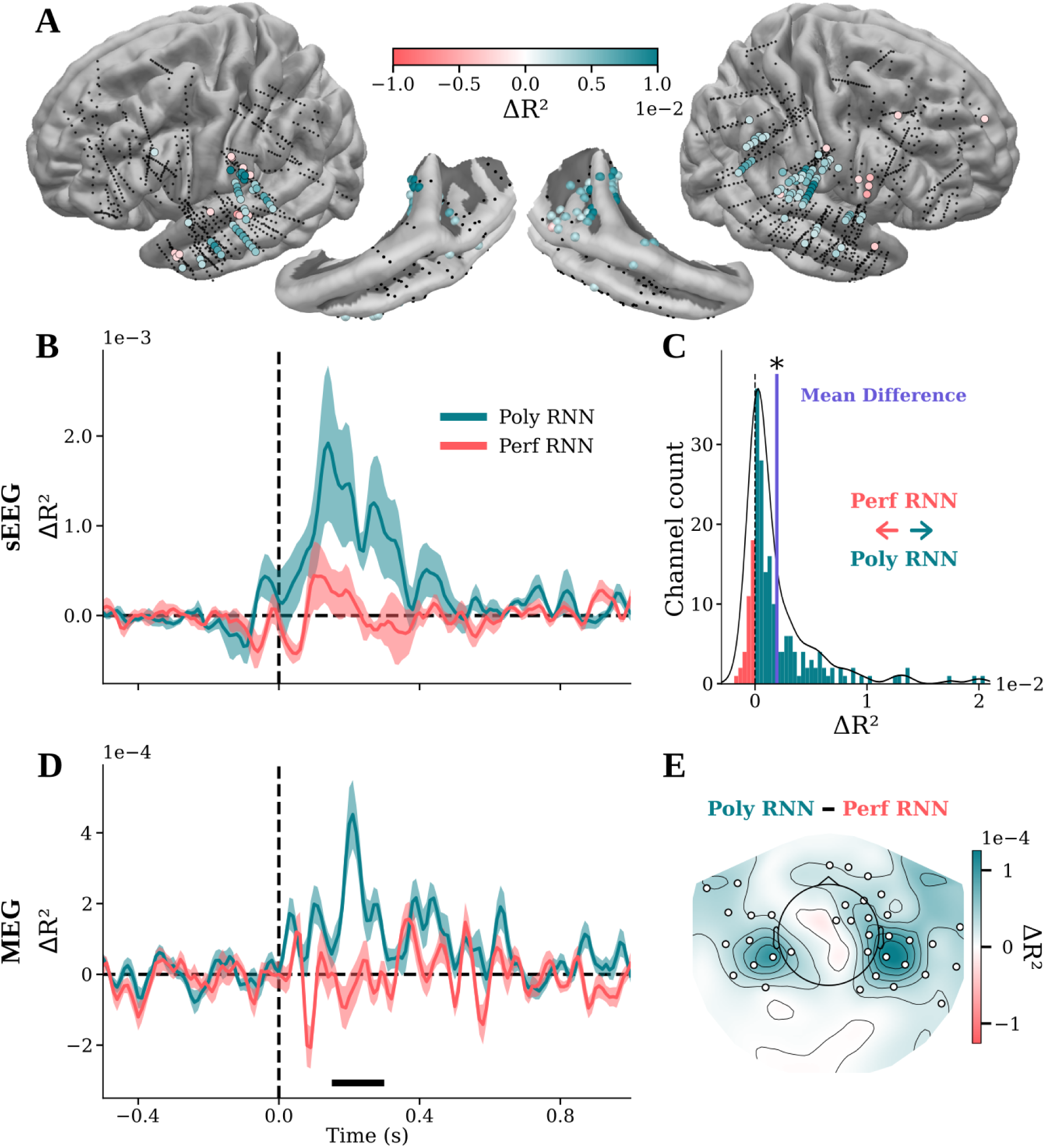
Comparison between PolyRNN and PerfRNN. (**A**) Spatial distribution of R² peak differences (within [-0.5, 1] s) between PolyRNN (cyan) and PerfRNN (red) in sEEG. Only channels with abs(ΔR²) > 0.002 are coloured, for visual purpose (no statistical testing). (**B**) Variance uniquely explained by each model, compared with control acoustic features, averaged across participants and auditory channels in sEEG. Shaded areas show SEM across channels. (**C**) Histogram of R² peak differences between PolyRNN and PerfRNN models in sEEG, across auditory channels. * t-test against zero, p < 0.001. (**D**) Same as (B) in MEG.The solid black ribbon indicates a significant difference between PolyRNN and PerfRNN (p < 0.05, cluster corrected). (**E**) Topography of R² difference between PolyRNN and PerfRNN at the peak (within [-0.5, 1] s), in MEG. Significant channels are indicated as white dots (p < 0.05, cluster corrected).

## Discussion

We have developed a novel recurrent neural network, PolyRNN, to track musical expectations during the listening of polyphonic pieces in a manner better aligned with human neurophysiological principles than previous state-of-the art models. PolyRNN takes advantage of the computational efficiency of neural networks to handle the complex structures occurring in natural music. It is trained on a large dataset of real recordings and functions with a piano roll representation to provide a plausible approximation of the learning and predictions of human listeners. Our analysis shows that PolyRNN makes better predictions in the musical genre it was trained on (classical music, compared to jazz or pop), suggesting that it captures musical structure relevant for a listener. Furthermore, the switch to a piano-roll format appears enough to markedly improve alignment of model predictions with human neural responses to music listening.

Specifically, PolyRNN successfully predicts neural activity in primary and associative auditory regions around 200 ms after notes onset. This effect is primarily driven by the encoding of surprise (prediction error) and is reminiscent of the event-related components such as the mismatch negativity and early-right anterior negativity found in seminal studies on musical expectations (Koelsch, 2009) and recent studies using monophonic stimuli (Omigie et al., 2019; Di Liberto et al., 2020; Kern et al., 2022; Shankaran et al., 2024). In addition, we show two other effects. First, surprise is also later encoded, peaking at 300 ms in sEEG and closer to 400 ms in MEG. These timings correspond to the P300 observed in other studies in response to harmonically surprising events in chord sequences (Janata, 1995; Regnault et al., 2001; Bigand et al., 2014), suggesting that PolyRNN’s surprisal is related to syntactic features encoded in the neural response. In MEG, this late component is also aligned with the 200 ms response to the next note, which could alternatively indicate that the response to the next note is modulated by surprise of the previous one. Second, the predicted density is also encoded in the neural data, but at a much earlier time-point. This suggests that predictions initially concern the mere occurrence of an event, regardless of its content, with pitch-related predictions emerging at a later stage. The model does not predict beyond the next timestep of the piano-roll (i.e. the next 50 ms), so the notion of “temporal prediction” as studied in music or cognitive science is not directly applicable (Rimmele et al., 2018; Zalta et al., 2024). However, predicted density likely represents the closest approximation in our model to assessing the likelihood of an event occurring at a given sample. Future work may focus on developing models capable of making predictions further into the future.

Our analysis shows that PolyRNN advances over the well-established IDyOM model in capturing neural responses, even for monophonic music (Di Liberto et al., 2020). We first replicated the encoding of IDyOM’s surprise with the EEG data from Di Liberto and colleagues (2022) and their set of control features (sound envelope and derivative), and found that PolyRNN best fit the neural response around 200 ms after note onset. We then added more control features representing low-order contextual regularities (pitch interval and time interval between consecutive notes), and PolyRNN retained its predictive value while IDyOM did not. This suggests that PolyRNN captures higher-order musical features encoded in brain activity. These features could correspond, for example, to the tonality or pitch range of the local context, and future projects may aim to identify them. Additionally, it is unclear what aspect of PolyRNN drives this neural prediction improvement, as both the training dataset (polyphonic music for PolyRNN, monophonic music for IDyOM), and the architecture (recurrent units for PolyRNN, Markov-chains for IDyOM) are different. From a training perspective, it could be that the model’s ability to process polyphony allows for use of a musical corpus that better reflects participant experience. On the other hand, it is possible that shifting to an RNN architecture enables better capture of higher-order relationships between notes that humans naturally perceive, even in single stream melodies. This would particularly support the growing trend in cognitive neurosciences to adopt RNN and Transformer models, which offer a more accurate representation of the neural encoding of perceptual statistical properties (Donhauser & Baillet, 2020; Saxe et al., 2021; Caucheteux & King, 2022; Kanwisher et al., 2023; Arana et al., 2024).

Another critical feature of PolyRNN is its musical representation and prediction task. While the generative music model Performance RNN (PerfRNN) is built with the same recurrent architecture and trained on the same data, it performs poorly in predicting neural responses to naturalistic, polyphonic music. In fact, PerfRNN processes simultaneous events in a serialized format (midi token), disrupting temporal organization of the stimulus in a way that doesn’t plausibly represent the perception and prediction of a human listener. In contrast, PolyRNN processes piano rolls and makes predictions for each possible pitch in parallel at each timestep of a sequence. One way to interpret a piano roll is to see it as a simplified cochleogram, a time-resolved sound representation known to emerge from the cochlea and to be conveyed via the auditory nerve to the auditory cortex (Chi et al., 2005). In other words, it constitutes a representational space that matches the computations taking place in the primary and associative auditory cortex. The superiority of PolyRNN at predicting the neural response to music thus highlights the importance of choosing adequate inputs and outputs for modeling the cognitive process of interest (Kanwisher et al., 2023).

Overall, our results validate the ability of PolyRNN to retrieve the neurophysiological markers of musical expectations in polyphonic music listening. This enables the use of naturalistic music in various paradigms, and brings the ecological validity that is crucial to study the perceptual, social and affective mechanisms involved in real-life music listening (Tervaniemi, 2023). This validation should be complemented with behavioral data in future work, as done with IDyOM (Pearce, 2005). Additionally, a promising perspective would be to explore the musical features captured by models like PolyRNN. While the black-box nature of neural networks make them difficult to interpret, methods have been developed to uncover the computations acquired through training (Olah et al., 2018). In the context of musical expectations, this could provide a way to explore how musical representations emerge depending on the musical background and on the constraints imposed by the prediction task.

## Materials and Method

### 1. Participants

#### 1.1. EEG

The open source EEG dataset from Di Liberto and colleagues (2020) was acquired on 20 healthy adults (10 females, mean age = 29 years). Half of them were musically trained, and the others had no musical background. They had no history of hearing impairment or neurological disorder.

#### 1.2. MEG

MEG signals were acquired on 27 normal-hearing participants (19 females, age = 28.3 ± 7.4 years) at the Institut du Cerveau, Paris, France. Musical experience was self-reported via the Goldsmiths Musical Sophistication Index (Müllensiefen et al., 2014). The General Sophistication value spanned from 26 to 102 (mean = 70, std = 18), showing a high diversity in musical experience across participants. This study was approved by the Comité de Protection des Personnes Tours OUEST 1 on 10/09/2020 (project identification number 2020T317 RIPH3 HPS).

#### 1.3. sEEG

10 patients (3 females, age = 28.1 ± 5.4 years) with pharmacoresistant epilepsy took part in the study. They were implanted with depth electrodes for clinical purposes at the Hôpital de la Timone (Marseille). Neural recordings were performed between 3 to 10 days after the implantation procedure. No sedation or analgesics drugs were used, and patients received their usual antiepileptic treatment. Recordings were always acquired more than 4 hours after the last seizure. Patients were included in the study if their implantation map covered at least partially the Heschl’s gyrus (left or right). Neuropsychological assessments carried out before SEEG recordings indicated that all patients met the criteria for normal hearing. None of them had their epileptogenic zone including the auditory areas as identified by experienced epileptologists. 3 patients have had musical training (> 5 years of instrumental practice), one being a professional musician. Informed consent was obtained from all patients. The study was approved by the Assistance Publique – Hôpitaux de Marseille (health data access portal registration number PADS E2YSEB). Recordings, interpretation and analysis of SEEG were performed following the French guidelines on stereoelectroencephalography (Isnard et al., 2018) and recommendations on SEEG analysis (Mercier et al., 2022).

### 2. Stimuli and procedure

#### 2.1. EEG

EEG data from Di Liberto and colleagues (2020) were acquired with ten excerpts of Bach pieces for solo instrument, with a duration of approximately 150s each. The audio stimuli were synthesized with piano sounds from the digital score. All stimuli were played 3 times throughout the experiment, for a total of 30 trials. Participants were asked to rate their familiarity to the piece at the end of each trial on a Likert scale. Behavioral data are not used here.

#### 2.2. MEG and sEEG

Stimuli for the two experiments were chosen from the MAESTRO database (Hawthorne et al, 2019), containing midi recordings of classical piano pieces (from baroque to modern eras) performed by skilled pianists. The midi files of the entire database were analyzed in 8 second clips with step size of 1 second. Within each 8-second window, the instantaneous note rate was assessed by identifying the timing of each note onset and taking the inverse of the difference between neighboring notes, discarding intervals less than 70 ms as part of chords.

After this, several features were recorded for each clip: Stability (inverse of the standard deviation of temporal intervals), Peak Frequency (most common note rate within bins of .5 Hz), Mean Frequency (mean note rate across frequencies), Onset Start (the time of the first note within the 8 second window) and Note Density (average number of concurrent notes played at each given point in time). From this information we selected the "best" clips as those with the highest Stability measurement within each Peak Frequency bin of 2.5 - 7.5 in 1 Hz steps. The 7.5 Hz condition was used only in the sEEG version of the experiment. Only clips with an average note density between 2 and 2.5 were considered. We avoided picking clips from the same songs by ensuring the levenshtein ratio between the titles and composers of each entry were not greater than .9. Selected clips were manually inspected to remove duplicates across multiple performances which could yield the same piece due to different naming practices. The 18 most stable clips were chosen to be included in the experiment.

Remaining clips were saved as individual midi files. They were then converted to sound files using the python package Pretty Midi (Raffel and Ellis, 2014) fluidsynth function. Midi files were snapped to note onset start to ensure no extra silence at the beginning of sound files). The soundfont used to generate the audio was a recording of a Mason Hamlin piano (MasonHamlin-A-v5.2.sf2) recorded and published online for free use at Soundfonts4u (https://sites.google.com/site/soundfonts4u). For a separate planned analysis, we were interested in altering the onset shapes of the notes and therefore used the original and several altered versions of the soundfont to smooth the piano’s acoustic edge. To do so, we used the polyphone soundfont editor to manipulate the attack in 3 settings: 0.001 seconds (original setting), 0.100 seconds (smooth setting), 0.250 seconds (smoothest setting), smoothing the onset shape of the piano sound. The onset manipulation was applied to specific clips separately so that participants would not hear the same clip with different onset shapes. The application was done in a balanced way across clips so that the mean stability rank for each onset shape was controlled. The sEEG data used all three onset shapes, while the MEG only used two (original and smoothest settings). The effect of onset shapes was not analyzed in the present study. Trial order was set and frozen across all participants. In the sEEG experiments, each clip was repeated twice while in the MEG version, clips were repeated 3 times. Trial order was balanced so that repetition numbers were as balanced as possible in terms of position within the experiment. Overall, the MEG and sEEG experiments contained 240 and 180 trials respectively.

Participants listened to 2 blocks of 90 stimuli in sEEG or 8 blocks of 30 stimuli in MEG. They completed an old/new task, where they had to indicate whether they already heard the same excerpt during the experiment. Their response was given with a button press at the end of each trial. Behavioral responses were not analyzed in the present study.

### 3. Acquisition and preprocessing

#### 3.1. EEG

EEG data from Di Liberto and colleagues (2020) were acquired with a 64-channels BioSemi Active Two System. To reproduce the preprocessing of the original paper, the signal was bandpass filtered (1 - 8 Hz) and downsampled (64 Hz) with MNE (v1.6) in a python environment. We re-epoched the data at the note level, keeping signals from -1 to 1 seconds relative to each note onset. Epochs based on notes occurring in the first 5 seconds of a trial were discarded to remove the initial auditory burst. Similarly, epochs centered on notes occurring in the last second of a trial were removed to avoid post-stimulus activity rebound. Overall, the data kept for analysis contained 18555 epochs per participant.

#### 3.2. MEG

MEG data was recorded with an Elekta Neuromag TRIUX system with a sampling frequency of 1000 Hz and a low-pass filter at 330 Hz. All the following preprocessing steps were performed in MNE-Python (Gramfort et al., 2013). Bad channels and signal portions were removed through automatic detection as well as visual inspection. We then used signal-space separation and Maxwell filtering to reduce external artifacts, compensate for head movement, and to reconstruct bad channels. A Notch filter at 50 Hz was applied to reduce electrical noise. Eye movements and cardiac electrical activity were filtered through Independent Component Analysis (ICA), using electrooculogram and electrocardiogram electrodes to automatically detect the corresponding components to discard. The 102 magnetometer channels were then selected, downsampled at 100 Hz and filtered between 0.5 and 12 Hz (zero-phase finite impulse response filter).

We re-epoched the data at the note level, keeping signals from -1 to 1 seconds relative to each note onset. Epochs based on notes occurring in the first 2 seconds of a trial were discarded to remove the initial auditory burst. Similarly, epochs centered on notes occurring in the last second of a trial were removed to avoid post-stimulus activity rebound. Finally, epochs based on notes played almost simultaneously (time interval < 50 ms) were removed, with only the first epoch of such a group being kept. Overall, the data kept for analysis contained a maximum of 12766 epochs per participant when no trials were removed during artifact rejection.

#### 3.3. sEEG

The sEEG signal was recorded using depth electrodes shafts of 0.8 mm diameter containing 10 to 15 electrode contacts (Dixi Medical or Alcis, Besançon, France). The contacts were 2 mm long and were spaced from each other by 1.5 mm. The locations of the electrode implantations were determined solely on clinical grounds. Patients were included in the study if their implantation map covered at least partially the Heschl’s gyrus (left or right). The cohort consists of 9 bilateral implantations and 1 unilateral implantation (right), yielding a total of 150 electrodes and 1690 contacts. Data were recorded using a 256-channels Natus amplifier (Deltamed system), sampled at 512 Hz and high-pass filtered at 0.16 Hz. A monopolar reference montage setup was used for the recording. The recording reference and ground were chosen by the clinical staff as two consecutive sEEG contacts on the same shaft both located in the white matter and/or at distance from any epileptic activity.

The precise localization of the channels was retrieved with a procedure similar to the one used in the iELVis toolbox (Groppe et al., 2017). First, we manually identified the location of each channel centroid on the post-implant CT scan using the Gardel software (Medina Villalon et al., 2018). Second, we performed volumetric segmentation and cortical reconstruction on the pre-implant MRI with the Freesurfer image analysis suite (documented and freely available for download online http://surfer.nmr.mgh.harvard.edu/). Third, the post-implant CT scan was coregistered to the pre-implant MRI via a rigid affine transformation and the pre-implant MRI was registered to the MNI template (MNI 152 Linear), via a linear and a non linear transformation from SPM12 methods (Penny et al., 2011), through the FieldTrip toolbox (Oostenveld et al., 2011). Based on the brain segmentation performed using SPM12 methods through the Fieldtrip toolbox, channels located outside of the brain were removed from the data (2.25 %).

The processing of continuous sEEG data was performed with MNE (v1.6) in a python environment. The signal was notch filtered at 50Hz and harmonics up to 250Hz to remove power line artifacts, bandpass filtered between 0.5 and 12 Hz and downsampled to 100Hz.

Channels and epochs with artifacts and epileptic spikes were discarded with a semi-automatic procedure (Mercier et al., 2022). sEEG data were epoched between -0.5 to 8.5 seconds relative to stimulus onset, and the peak of the absolute value of the signal was taken for each epoch of each channel. For channel rejection, peak values were first averaged across epochs. Then, the mean and the standard deviations of peak values across channels were computed, and a threshold was defined as N standard deviations above the mean. The factor N was set based on visual inspection of the data (in a range from 4 to 10), and all channels with a peak value exceeding the threshold were removed from subsequent analyses. For epoch rejection, the same procedure was applied with peak values averaged across channels. This whole procedure was conducted for each patient independently, removing a total of 3 channels and 1 epochs. Additionally, a technical issue made the triggers unusable for half of the data of 1 patient. These data have been removed from all analyses.

The sEEG data was then re-epoched at the note level, following the procedure used in MEG, yielding 10795 events when no trials were rejected.

### 4. Models of musical expectancies

#### 4.1. IDyOM

The Information Dynamics Of Music (IDyOM; Pearce, 2005) model was designed to estimate the conditional probability of the pitch and onset time of upcoming notes in a musical sequence. IDyOM features a long-term memory (LTM; trained on a corpus) and a short-term memory (STM; capturing the regularities in the current piece). IDyOM is limited to the processing of sequences of single events, and has mainly been used with monophonic music (Omigie et al., 2012; 2019; Gold et al., 2019; Quiroga-Martinez et al., 2019; 2020; Di Liberto et al., 2020; Kern et al., 2022) or chord sequences (Cheung et al., 2019; 2024), and cannot process polyphonic pieces. Similarly to Di Liberto and colleagues (2020), we trained IDyOM on a large corpus of Western tonal music, including Canadian folk songs (Creighton, 1966), German folk songs from the Essen folk song collection (Schaffrath, 1992), and chorale melodies harmonized by Bach (Riemenschneider, 1941) as in other applications of the model (e.g., Pearce, 2005; Pearce and Wiggins, 2006; Egermann et al., 2013; Hansen and Pearce, 2014; Gold et al, 2019). This corpus did not contain the stimuli used by Di Liberto and colleagues. IDyOM was configured with STM and LTM combined, and was trained with the *pitch* and *note onset* (i.e. timing) viewpoints separately. For each note of each stimulus, we retrieved probability distributions of pitch and timing, and computed the *surprise* (Equation 1) and *uncertainty* (Equation 2).

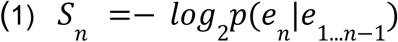

where *p*(*e_n_*|*e*_1…*n*−1_) is the conditional probability of the pitch or timing of the *n^th^* event of the sequence.

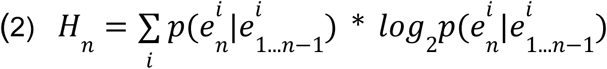

#### 4.2. Performance RNN

Performance RNN (PerfRNN) is a recurrent neural network model designed to generate piano music. It is based on a musical representation analogous to digital scores (MIDI format), with discrete events of different types: note onsets (pitch), note offsets (pitch), and time intervals between events. In every processing step, PerfRNN generates a probability distribution over all possible event types and values and pulls one specific event out before starting the process again. Polyphony is thus handled as a serialized process, with simultaneous notes generated one after the other with no time interval in between. The model can also be applied to existing music, generating probability distributions for every event of our stimuli. We used an available version of PerfRNN (http://download.magenta.tensorflow.org/models/performance.mag) that was pre-trained on real performances from the International Piano-e-Competition dataset (renamed MAESTRO at time of publication; https://magenta.tensorflow.org/datasets/maestro). It is a recurrent neural network with 3 long short-term memory (LSTM) layers (see Oore et al., 2020 for details). For each note of each stimulus, we retrieved probability distributions of note onset and time interval events separately, and computed the *surprise* (Equation 1) and *uncertainty* (Equation 2). When multiple notes start at the same timing, they were all assigned the mean surprise and uncertainty value across them, and duplicate events were removed later in the analysis pipeline (see Section Acquisition and preprocessing).

#### 4.3. PolyRNN

We designed a recurrent neural network, PolyRNN, to process time-resolved, polyphonic musical sequences (https://github.com/pl-robert/musecog). The model takes as input a sequence of timesteps, with a vector representing the onset, sustain or absence of each possible note at each timestep (a “piano-roll”). PolyRNN is trained to perform a multi-item prediction task, where a probability of occurrence is estimated for each note of the next timestep (e.g. the model could output a probability of 1 for several notes, or 0 for all notes). This choice of representation implies that the model doesn’t make any prediction about the velocity and the duration of upcoming notes, making it unable to generate music. Furthermore, this representation does not feature an explicit stream segregation. Even if the model learns to separate different streams (e.g. a melody line and a bass line), it will not appear explicitly in its output.

PolyRNN was trained on the International Piano-e-Competition dataset (renamed MAESTRO at time of publication; https://magenta.tensorflow.org/datasets/maestro), containing approximately 150 hours of classical piano music (from XVII^th^ to XX^th^ centuries) performed by professional pianists and recorded in the MIDI format. MIDI files were converted to piano rolls (2D arrays with dimensions n_pitch and n_timesteps) with a frequency sampling of 20 Hz. All velocity information was removed, the onset of a note was coded as a 2, a sustained note as a 1, and an absence of note as a 0. To avoid any bias toward the most common scales in classical music, the pitch of all notes of a given piece were randomly raised or lowered by +/- 6 semitones during training. The dataset was randomly divided in training (80%) and validation (20%) sets.

PolyRNN was built with the pytorch python package (Paszke et al., 2019). The model contains a single Long-Short Term Memory (LSTM) layer (input size = 88, hidden size = 88, dropout = 0.05) connected to a linear layer (output size = 88) with a sigmoid activation function. PolyRNN was trained with batch learning (batch size = 15, batch length = 60 sec), truncated back-propagation through time (TBPTT, window = 5 sec) and an Adam optimizer (learning rate = 10^-5^). The model’s performance was measured with binary cross entropy (log base 2). The training was manually stopped at batch number 100K, after training and validation losses had stagnated for the last 20K batches (Fig. S6). The final training loss and validation loss averaged across the last 1K batches were 3.41 and 3.49 bits/timestep respectively. Other versions of the model have been trained with LSTM layers ranging from 1 to 5, LSTM hidden size from 88 to 512, batch size from 4 to 32, TBPTT from 2 to 10 seconds and learning rates from 10^-5^ to 10^-2^, but didn’t reach a lower validation loss.

Qualitatively, visual inspection of the output of PolyRNN suggests that its prediction dynamically adapts to the note rate, the pitch range and the tonality of the preceding musical context (Video S1). It further makes timing predictions when the input keeps a steady rhythm, and predicts simple pitch trajectories like ascending or descending scales, arpeggios, and trills. Consistent with the known limitations of recurrent neural networks (Williams & Peng, 1990), the model seems to forget the events that happened past the TBPTT window, and is unable to catch precise patterns (e.g., specific themes or broken chords) or long-term structure (e.g., second presentation of a theme). Overall, PolyRNN seems to perform the non-trivial task of dynamically retrieving multiple relevant musical features (note rate, pitch range, tonality, pitch trajectories…) and represents a plausible implementation of predictive processing derived from statistical learning.

Quantitatively, the predictions of PolyRNN cannot be directly evaluated through benchmarking and model comparison, as its prediction task is not comparable to those of existing music models. Instead, we compared the model’s performance on new piano datasets with musical genres matching (classical) or differing (jazz, pop) from the training corpus, as a means to test whether PolyRNN captured regularities specific to its training genre. We collected natural piano recordings (WAV format) from public playlists available on Spotify in the three musical genres, and we obtained 502 classical pieces (total duration of 34 hours, mean duration of 245 s, SD = 167 s), 823 jazz pieces (total duration of 69 hours, mean duration of 301 s, SD = 221 s) and 319 pop pieces (total duration of 19 hours, mean duration of 212 s, SD = 55 s) for a total of 1644 pieces, all of which are played by one piano only (mono-instrument). We then converted the audio files into MIDI files with the Onsets and Frames Transcription tool (Hawthorne et al., 2017). To evaluate PolyRNN’s performance between genres, we compared the binary cross entropy at note onset averaged within pieces with Welch’s t-tests (two-sided) and Bonferroni correction for multiple comparison (Fig. 1B).

The stimuli (see Section Stimuli) were then processed with PolyRNN, and three features of interest were derived from its predictions. First, the *surprise* value S_t_ of a note was computed as the binary cross entropy (same as the loss function used during training) of the timestep where the note appears (Equation 3). When multiple notes appear at the same timestep, they were all given the same surprise value, and duplicate events were removed later in the analysis pipeline (see Section Acquisition and preprocessing). Second, the *uncertainty* value of a note H_t_ was obtained by normalizing the model’s output of the timestep where the note appears and computing its entropy (Equation 4). This feature was chosen to allow for a direct comparison with the uncertainty derived from IDyOM in previous studies (Cheung et al., 2019; Gold et al., 2020; Di Liberto et al., 2020). However, its application in this modeling approach implies that it will be affected by the number of simultaneous notes happening in the current context (i.e. predicting 10 different notes will lead to a higher uncertainty than predicting only 1 note, even if it is a correct prediction). Finally, the *predicted density* D_t_ value of a note was computed as the sum of all the probabilities at the timestep where the note appears (Equation 5). In contrast with other models, PolyRNN is predicting the occurrence of each note (with their own probability between 0 and 1) at each timestep. Thus, the predicted density is expected to vary depending on the beat (i.e. high on the beats, low between the beats) and on the number of predicted notes.

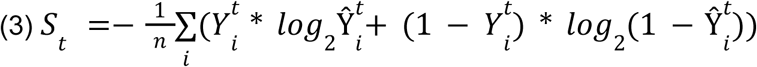

where Ŷ^*t*^ is the vector of occurrence probability estimated by PolyRNN at a time step *t*, and *Y*^*t*^ is the ground truth with a note onset coded as 1 and 0 otherwise.

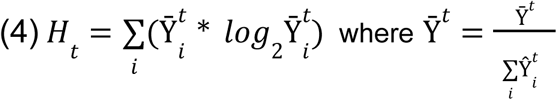

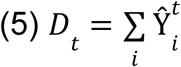

### 5. Control features

A set of control features was selected based on note properties (from original MIDI files) and acoustic features (derived from audio stimuli). Note properties contained the absolute time (since trial start), time interval (since last note), note rate (constant within a trial), absolute pitch (in log scale), and pitch interval (since last note). When multiple notes were played simultaneously, only the pitch of the highest note was kept.

In EEG analyses, acoustic features only contained the broadband amplitude envelope (Hilbert transform) and its half-way rectified derivative. These features were included in the available dataset but not the audio files of the stimuli, preventing us from computing other acoustic descriptors.

In MEG and sEEG analyses, acoustic features contained several time-varying descriptors of auditory sensations (Giordano et al., 2023), such as loudness, periodicity, timbre brightness and roughness (temporal resolution = 1 ms). Time-varying loudness and brightness were derived from the instantaneous specific loudness of the input signal (Glasberg & Moore, 2002), as estimated in the Genesis Loudness Toolbox in MATLAB. Instantaneous specific loudness measures the time-varying contribution to the overall instantaneous loudness (temporal resolution = 1 ms) in separate frequency bands (N frequency bands = 153, equally spaced on an ERB-rate scale between 20 and 15,000 Hz). For each temporal frame, loudness (measured on a sone scale) was then defined as the sum of the instantaneous specific loudness across frequency bands, whereas timbre brightness was defined as the spectral centroid, that is, as the specific loudness weighted average ERB-rate frequency (McAdams et al., 1995). Time-varying periodicity (ratio of periodic to aperiodic power, in dB) were estimated using the Yin model (Cheveigné & Kawahara, 2002). Time-varying roughness (Vencovský, 2016) was estimated using the model implemented in the MIRtoolbox v.1.7.2 in MATLAB. Spectral flux (rate of timbral change) was added as a fifth acoustic feature, as it has been shown to be strongly encoded in the brain’s response to music (Weineck et al., 2022). Spectral flux was estimated with the Surfboard python package v.0.2.0 (window size = 0.04 s, hop size = 0.01 s). The derivatives of each acoustic feature were also added to the set of control features, except in the case of loudness, where the half-way rectified derivative was used. Finally, all acoustic features were averaged in a 150 ms time-window starting from note onsets, to obtain a single value per epoch.

### 6. Encoding analysis of PolyRNN music features

In all datasets, a multiple linear ridge regression was used to model the amplitude of the brain response to individual notes (or groups of simultaneous notes). For each channel and each time-point (from -1 to 1 seconds; see Section Acquisition and preprocessing) of the response separately, the linear model was fitted and tested following a 5-fold cross-validation scheme, yielding a measure of predicted variance (cv-R²). This was repeated with several ridge parameters (alphas ranging from 10^2^ to 10^7^), and the best cv-R² were kept. This flat cross-validation setup was preferred over a nested cross-validation scheme, to reduce the computational cost (see below, permutation tests) and because of similar overall quality between both procedures when only a few hyperparameters (here n=1) have to be optimized (Wainer & Cawley, 2021). Importantly, the same procedure was applied for the different linear models of interest and the permuted analyses (see below).

To assess the specific encoding of musical expectancies, we focused on the variance uniquely explained by music models features. It was retrieved by computing the cv-R² from all acoustic control features (model A), or from the full set of control and music model features together (model AM), and taking the difference (ΔR² = cv-R²_AM_ - cv-R²_A_).

In our ecological stimuli, the time interval between consecutive notes is short (mean time interval = 197 ± 77ms) and is likely to induce overlapping neural responses. Since the music models features are not independent across consecutive events (positive autocorrelation; Fig. S7), the values associated with the current note *n* may partly predict the neural response to the following note *n + 1*. To allow the current analysis to disentangle late components in response to *n* from early components to *n + 1*, we computed the regression with the features associated with both notes (*n* and *n + 1*), and we separated the contribution of each with variance partitioning (later in the analysis pipeline; see below). In all datasets, the final linear model A thus contains all acoustic control features of notes n and *n + 1*. The final AM_PolyRNN_ model contains the same features as A, with the addition of *Surprise*, *Uncertainty*, *Predicted density* of notes *n and n + 1*.

In MEG and sEEG, statistical significance was assessed with cluster level permutation tests. The regression model AM_PolyRNN_ was estimated 1000 times with random permutations of the music model features’ values across epochs. Chance-level ΔR² values were computed by taking the difference in cv-R² between model A and the permuted versions of AM_PolyRNN_.

In MEG, group-level statistics were obtained with a one-tailed t-test (ΔR² > 0) on both original and permuted results. Spatio-temporal clusters were defined with MNE’s _find_clusters function, in which channels and time points with a t-statistic that exceeds an arbitrary threshold (t-threshold = 2.63) are clustered based on adjacency in time (consecutive time points) and space (adjacent channels on the MEG topography using MNE’s find_ch_adjacency function). The null-distribution was obtained by taking the cluster of highest magnitude (mean t-statistic) in each permutation. In the original data, a cluster is significant when its magnitude exceeds at least 95% of the cluster magnitudes of the null distribution.

In MEG, all the channels belonging to at least one significant spatio-temporal cluster were separated into two spatial groups (left and right). A sign-based permutation temporal cluster test (permutation_cluster_1samp_test in MNE) on the spatial averages was used to test for lateralization effects (10000 permutations, t-threshold = 2.06, two-tailed).

In sEEG, the heterogeneity of channel locations prevents us from comparing channels across participants and computing group-level statistics. Hence, clusters were based on ΔR² values (ΔR² threshold = 0.005) and temporal adjacency between data points. The null-distribution and statistical threshold were obtained in a similar way as with MEG data.

To complement these analyses with an independent channel selection, we define the auditory channels as the 20 channels with the highest M100 amplitude for each participant in MEG, and the 20 channels with highest magnitude (absolute value of amplitude) at 100ms after note onset per participant in sEEG. We computed the average of peak R² values (within [-0.5, 1] s) across these channels, and compared the linear models A and AM_PolyRNN_ with two-tailed paired t-tests.

Finally, we performed a variance partitioning analysis to evaluate the contribution of each music model feature in significant clusters. In AM_PolyRNN_, the unique variance of a given feature was obtained by taking the difference in cv-R² values between full AM_PolyRNN_ and AM_PolyRNN_ without that feature.The shared variance (between at least 2 variables) was defined as the cv-R² of the full AM_PolyRNN_ minus the sum of all unique variances.

### 7. Model comparison

#### 7.1. PolyRNN vs IDyOM

In EEG, the encoding of IDyOM’s expectations were estimated with a linear model AM_Idyom_ (five-fold cross-validated ridge regression) containing IDyOM’s *Pitch-Surprise*, *Pitch-Uncertainty*, *Timing-Surprise*, *Timing-Uncertainty* of notes n and n + 1. The acoustic control model A contained the sound envelope, its derivative, and all note properties (note rate, absolute time, time interval, absolute pitch, pitch interval) as regressors. To allow for a direct comparison with the results of Di Liberto and colleagues (2020), we also estimated a model A_env_ with the envelope and its derivative only, as well as full models A_env_M_PolyRNN_ and A_env_M_Idyom_ without any shifted feature (i.e., of note n + 1).

We first retrieved the spatio-temporal extent of the main effect obtained with A_env_M_Idyom_ - A_env_. Using an arbitrary threshold (ΔR² > 0.00035), we obtained a mask covering central electrodes and timelags ranging from 156 to 220 ms. For each linear model applied on EEG data, we took the average of ΔR² values within this mask, and compared them with 0 with one-tailed t-tests and Bonferroni correction. To compare the music models, we contrasted PolyRNN and IDyOM with either the envelope-only control (A_env_M_PolyRNN_ - A_env_ versus A_env_M_Idyom_ - A_env_) or the full acoustic control (AM_PolyRNN_ - A versus AM_Idyom_ - A) with two-tailed paired t-tests and Bonferroni correction.

#### 7.2. PolyRNN vs PerfRNN

As with PolyRNN, a multiple linear ridge regression was used to model the amplitude of the brain response to individual notes in both sEEG and MEG. The AM_PerfRNN_ model contains the same features as A (see Section Control features), plus *Pitch-Surprise*, *Pitch-Uncertainty*, *Timing-Surprise*, *Timing-Uncertainty* of notes n and n + 1. PolyRNN and PerfRNN were explicitly compared by taking the difference between cv-R²_PolyRNN_ and cv-R²_PerfRNN_ for each timepoint of each channel.

In MEG, statistical significance was assessed with sign-based permutation tests at the spatio-temporal cluster level as implemented in MNE’s spatio_temporal_cluster_1samp_test function (10000 permutations, T threshold = 2.53, two-tailed).

In sEEG, the difference between the peak-R²_PolyRNN_ and the peak-R²_PerfRNN_ was computed for each channel. Statistical significance was assessed with a two-tailed t-test across the auditory channels, defined as the 20 channels with highest magnitude (absolute value of amplitude) at 100ms after note onset per participant in sEEG (i.e. 200 channels).

## Acknowledgments

We thank Bruno Giordano for his support with data analysis. This research is co-funded by the European Union (ERC, SPEEDY, ERC-CoG-101043344), supported by Fondation Pour l’Audition (FPA RD-2022-09; FPA RD-2020-10), the Fondation Fyssen, grants from France 2030 (ANR-24-CE17-7152-03; ANR-16-CONV-0002) and the Excellence Initiative of Aix-Marseille University (A*MIDEX).

## Supplementary materials

https://www.youtube.com/watch?v=WTHKQMljzXY

**Video S1. Demonstration of PolyRNN’s predictions.**

**Figure S1.**
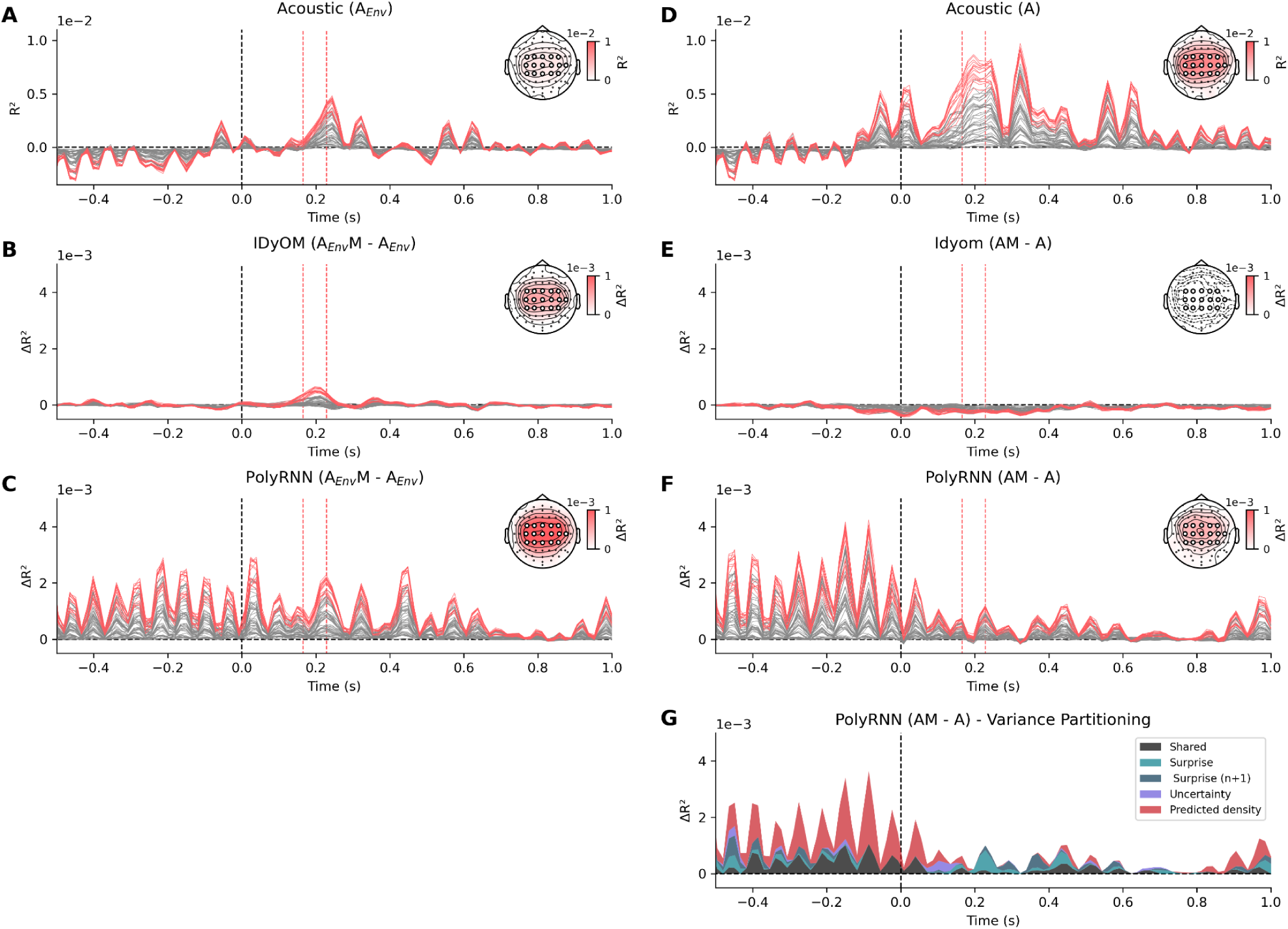
Encoding of acoustic and models (IDyOM and& PolyRNN) features in EEG. **(A)** Variance explained by control acoustic features (envelope and its half-way rectified derivative; A_Env_) in the data from Di Liberto and colleagues (2020). Red lines indicate the central channels selected in Figure 2. **(B)** Variance uniquely explained by IDyOM’s features (ΔR² = A_Env_M - A_Env_). This result mirrors that of Di Liberto and Colleagues. **(C)** Variance uniquely explained by PolyRNN’s features (ΔR² = A_Env_M - A_Env_). **(D-E-F)** Same as (A-B-C), but using the full set of control features (A; see Method). **(G)** Cumulative plot of variance explained, partitioned by each PolyRNN music feature (ΔR² = AM_All_ - AM_All-1_), and averaged across selected channels. Surprise (n+1) corresponds to the surprise of the next note. The shared variance represents all non-unique explained variances. The Y-axis is depicted from zero as negative cross-validated R2 is meaningless.

**Figure S2.**
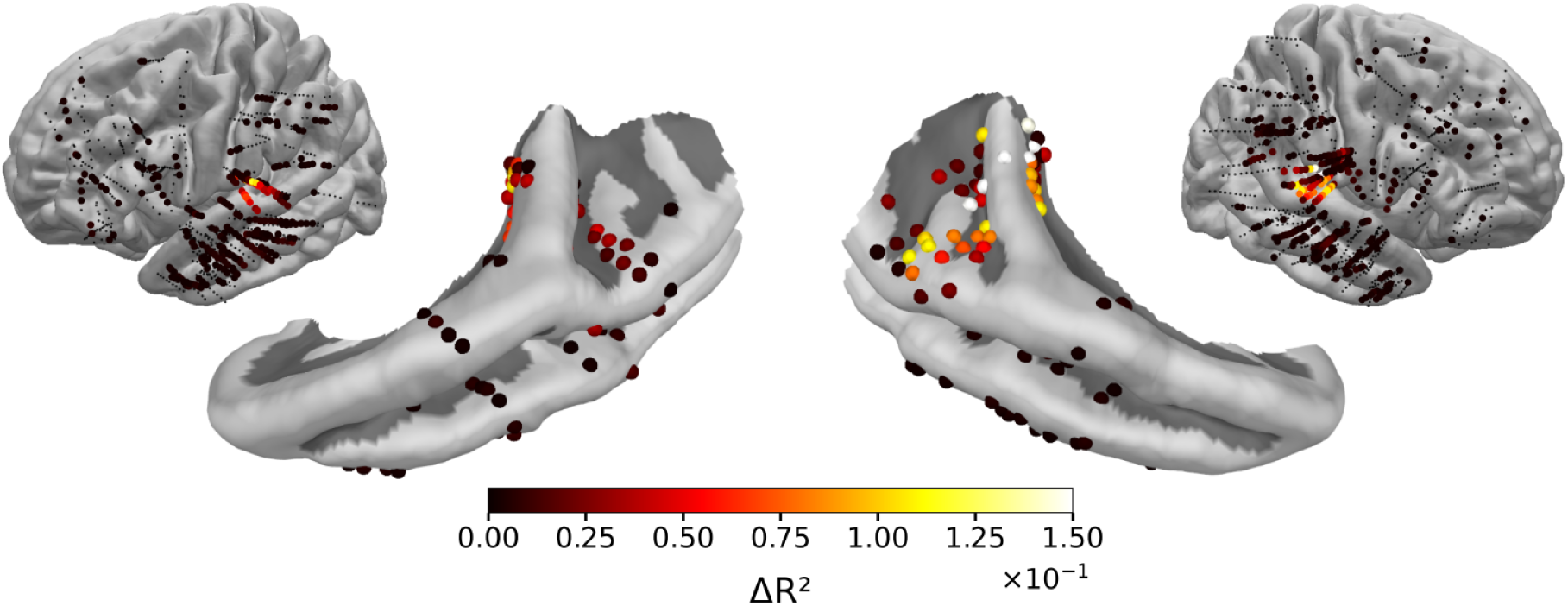
Encoding of acoustic features in sEEG. Variance explained by acoustic features, plotted on whole brains and temporal lobes. Peak R² (within [-0.5, 1] s) are color coded in significant sEEG channels. Only channels with abs(ΔR²) > 0.002 are coloured, for visual purpose.

**Figure S3.**
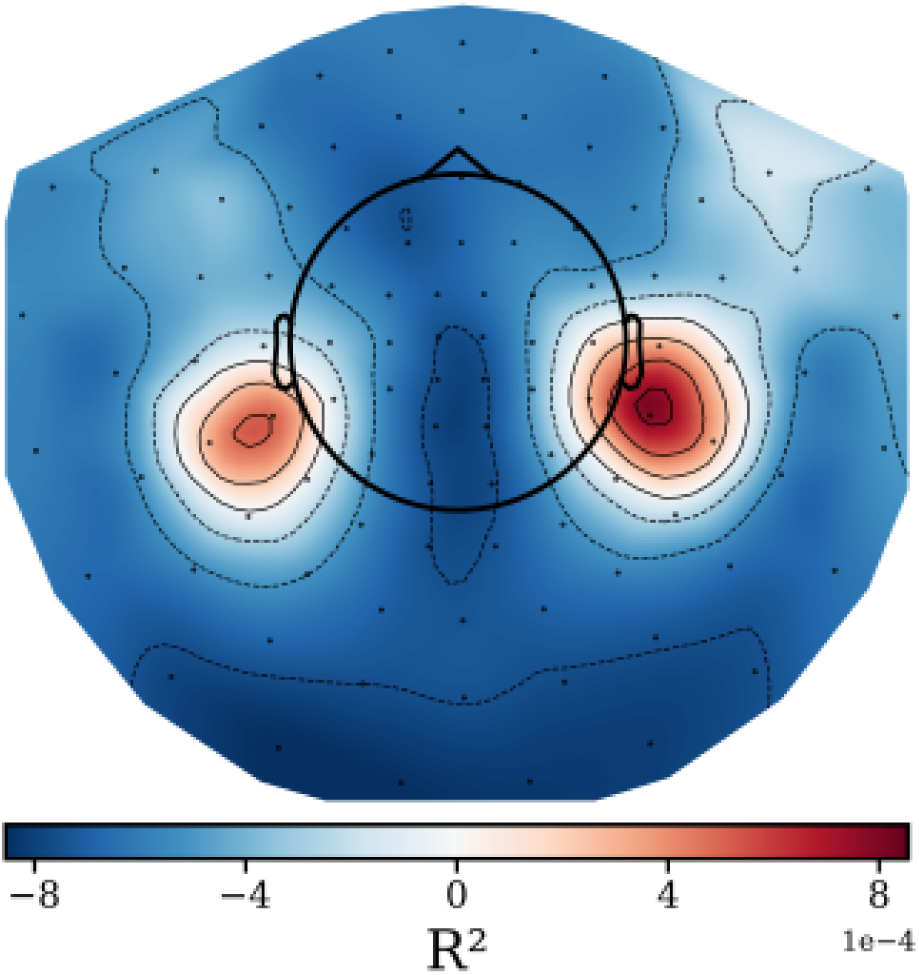
Encoding of acoustic features in MEG. Topography of variance explained by acoustic features, averaged across time ([-0.5, 1] s) and participants.

**Figure S4.**
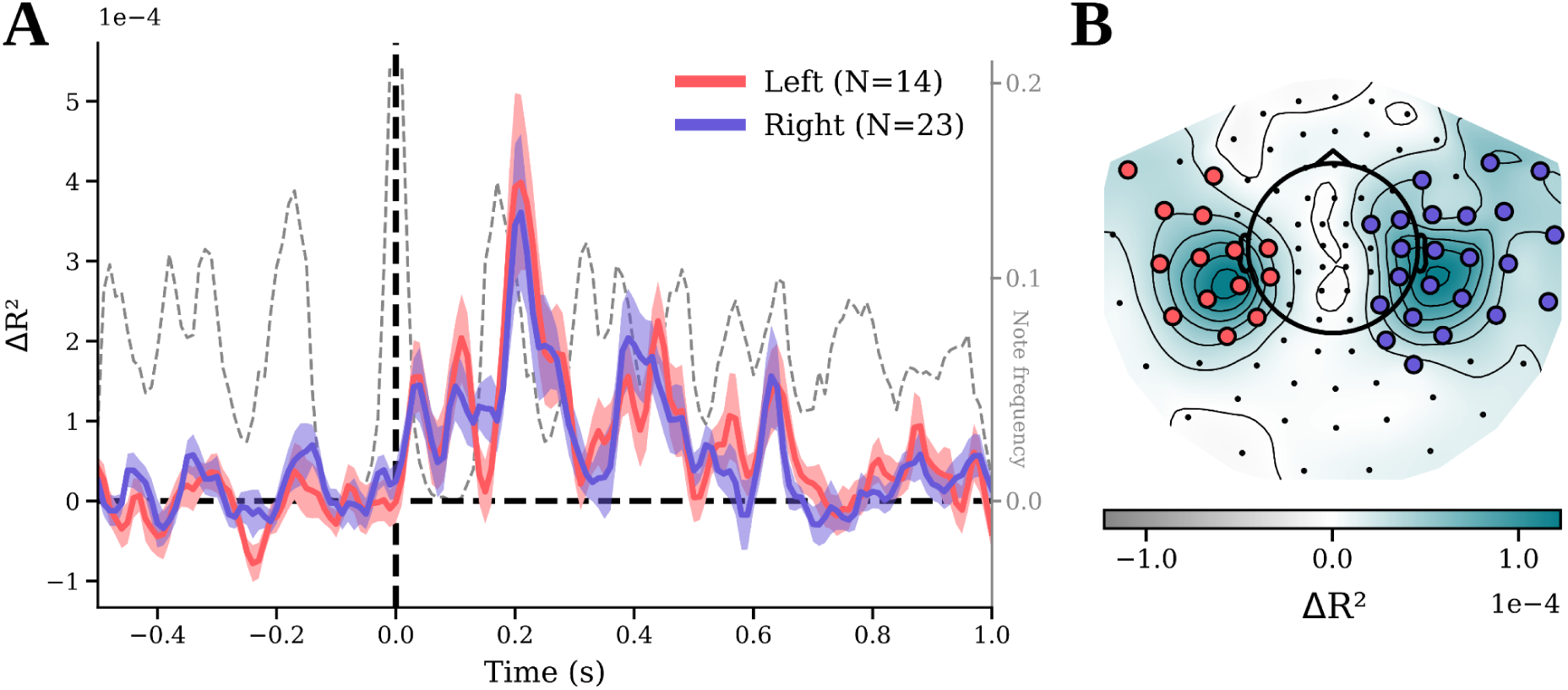
Encoding of polyphonic expectations (PolyRNN) in left and right hemispheres, in MEG. **(A)** Variance explained by PolyRNN features at the note-level (note onsets locked at t = 0s) in MEG, separated by hemisphere in significant channels. Shaded areas are SEM across channels. Note frequency represents the frequency of occurrence of a note at each timepoint (p = 1 at t = 0s, right y-axis). **(B)** Topography of ΔR² values averaged across time ([-0.5, 1] s) and participants in MEG data (N = 27). Significant channels in each hemisphere are indicated as colored dots (red: left, blue: right).

**Figure S5.**
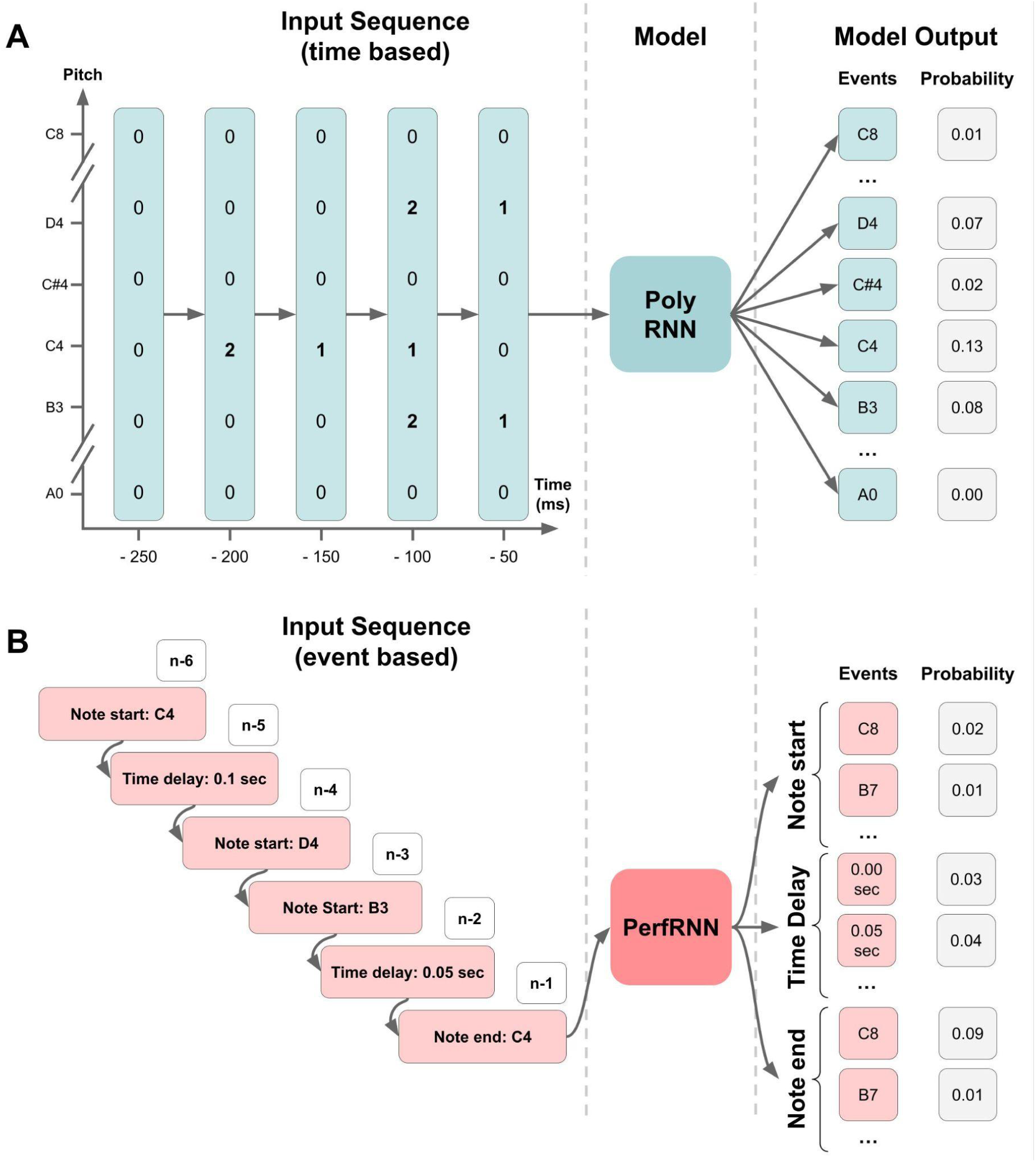
Comparison of PolyRNN and PerfRNN working principles. **(A)** PolyRNN receives sequences of vectors coding for the start (2), the sustain (1) or the absence (0) of each possible note of the 88 notes of the piano keyboard, with one vector per timestep (50 ms). With this representation, multiple note onsets can be represented within a single timestep. The model outputs predictions about the next timestep with one probability of occurrence per note. **(B)** PerfRNN receives sequences of events representing the pitch of a new note (note start), the length of a time interval (time delay) or the pitch of a note that ends (note end). Internally, these events are represented with a one-hot encoding. This representation is analogous to the event sequence contained in a MIDI file. The model outputs predictions about the next element in the sequence, with one probability for each possible event. When PerfRNN is used for music generation, the most probable event is chosen with a softmax function, and is added to the sequence.

**Figure S6.**
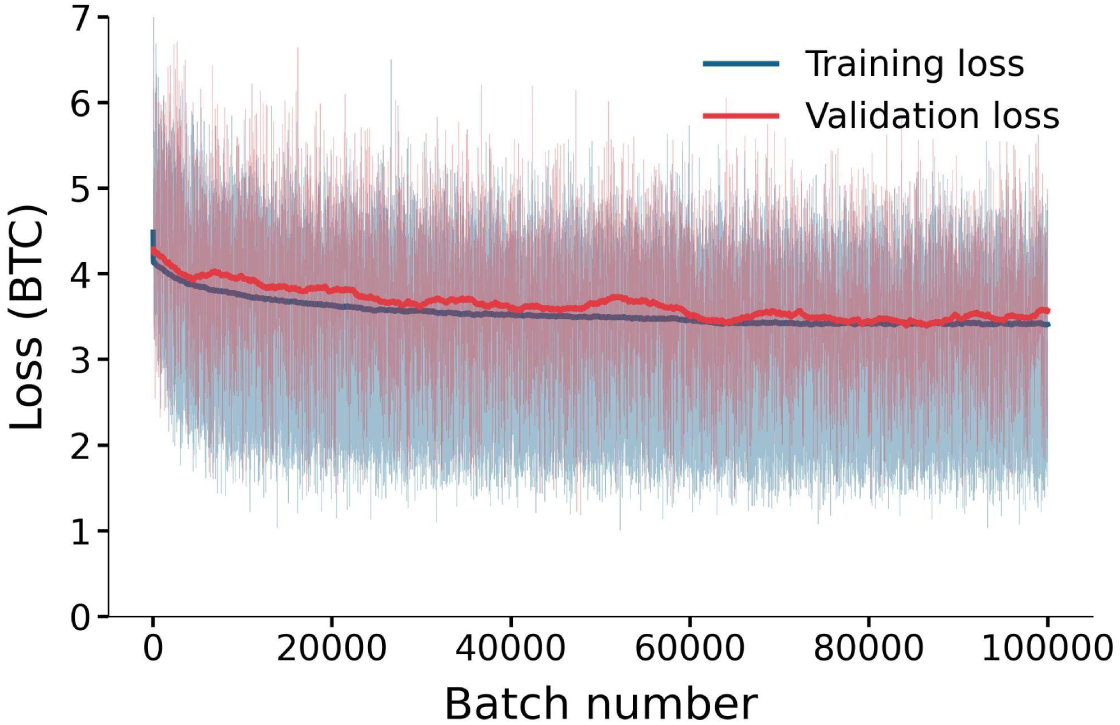
Training and validation loss of PolyRNN.

**Figure S7.**
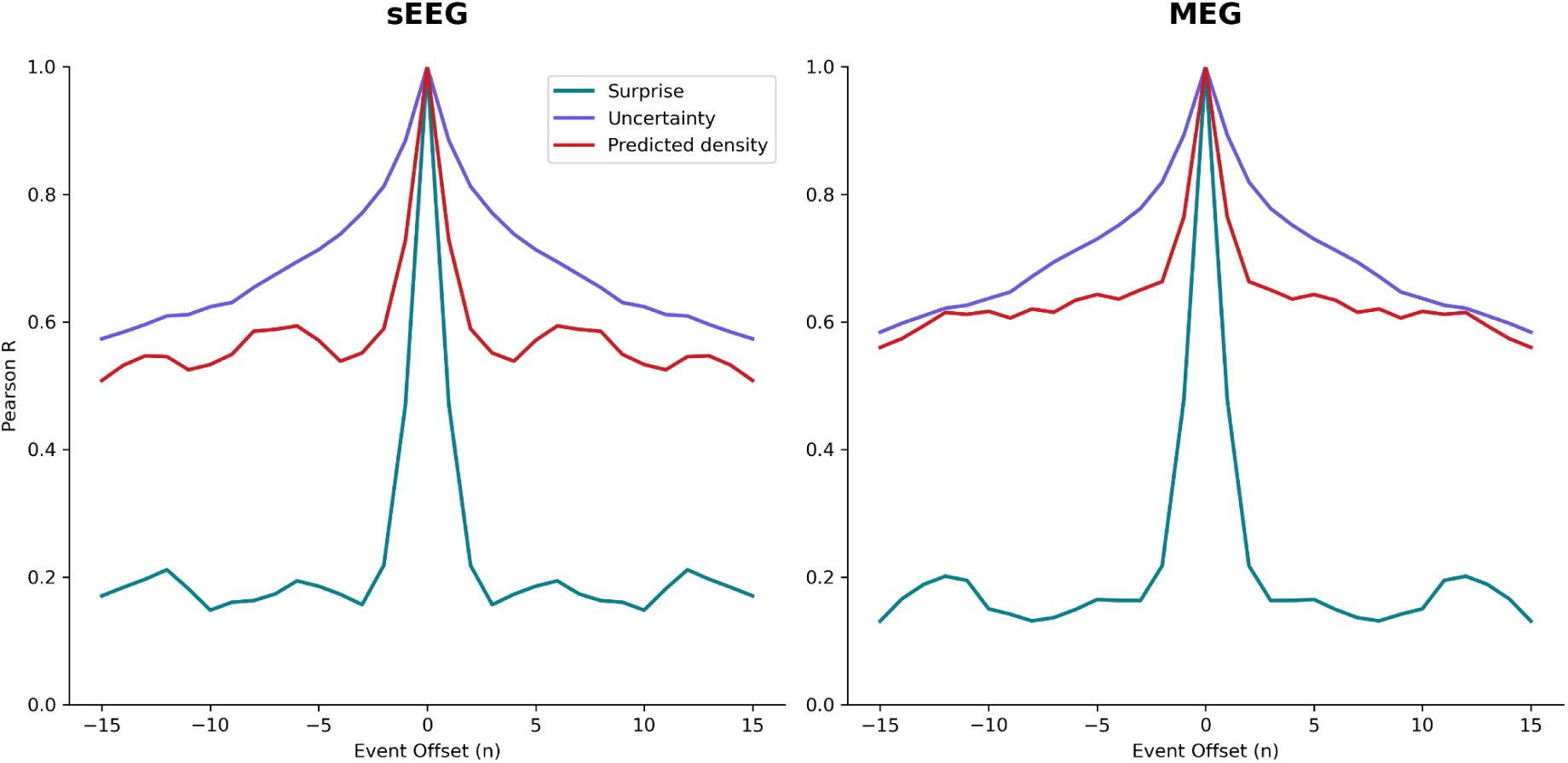
Autocorrelation of PolyRNN features. The autocorrelation is computed across the consecutive events (e.g. notes or groups of notes, see Method) in the stimuli used for sEEG and MEG data acquisition.

